# Omega-3 fatty acid normalizes postsynaptic density proteins via miRNAs regulation in hippocampus and prevents DEHP-induced impairment of learning and memory in mice

**DOI:** 10.1101/2024.10.23.619821

**Authors:** Muyao Ding, Hongyu Ma, Hui Du, Yinglong Yang, Min Yu, Cong Zhang

## Abstract

DEHP is the most widely used plasticizer in many products. However, growing evidence has indicated that DEHP may induce neurotoxicity. DEHP exposure affects mircoRNAs (miRNAs) expression in brain. A growing body of evidence suggests that nutrients and other bioactive food components prevent neurotoxicity through regulation of miRNA expression. Due to the increasing concern about the risks of DEHP to human health, we explored the neuroprotective effect of Omega-3 fatty acid (Omega-3FA) on subchronic DEHP-induced neurotoxicity in mice, and the potential involved miRNAs and their targets in the protective action of Omega-3FA against DEHP-induced neurotoxicity. Omega-3FA protected against the DEHP-induced impairment of learning and memory and alleviated the thinning of postsynaptic density (PSD) thickness in hippocampal synapses. We observed that there are fourteen up or down regulated miRNAs associated PSD in DEHP exposure which were normalized by Omega-3FA treatment. Protein targets in PSD of these differentially expressed miRNAs were predicted. Furthermore, the expression levels of protein mGluR5, Homer1, and NMDAR2B were carried out via Western blot, for further verifying PSD associated miRNAs’ targets are involved in neuroprotection of Omega-3FA against DEHP. These findings suggested that Omega-3FA protected DEHP-induced impairment of learning and memory as well as synaptic structure alteration in the hippocampus by regulating the expression of PSD associated miRNAs and their targets. Thus, Omega-3FA should be included in diet to prevent or suppress neurotoxicity caused by continuous exposure to DEHP.

## Introduction

Di-ethyl-hexyl-phthalate (DEHP) is widely used as plasticizer and softener in many consumer products, including building materials, food packaging, children’s toys, medical devices, and cosmetics [1]. DEHP is not chemically bound to the end products, it may easily transfer into the environment and then enter the body through ingestion, inhalation and skin exposure [2, 3]. Epidemiological studies have demonstrated that developmental DEHP exposure affected the neurodevelopment of newborns and decreased the intelligence of children [4, 5]. Defects in learning tasks and changes in behavior were observed in animals exposed to DEHP [6]. In summary, these findings suggest that DEHP causes neurotoxic effects in the central nervous system, including learning and memory impairment.

Synaptic plasticity in the hippocampus is considered to be the basis of learning and memory. Changes in synaptic functional plasticity are tightly associated with alterations of synaptic structure [7, 8]. Postsynaptic density (PSD) is a protein-dense complex attached to the postsynaptic membranes, contains membrane receptors, scaffold proteins, kinases and many signaling molecules and is crucial in synaptic signal transduction [8, 9]. The change in PSD thickness serve as an important indicator which reflects postsynaptic structural changes in hippocampus-dependent learning [10]. Moreover, PSD protein content is involved in a long-term increase in the synaptic strength of LTP [11]. Recent studies have reported that DEHP downregulated the expression of postsynaptic density protein 95 (PSD 95) and synapsin-1 [12], which play crucial roles in synaptic structure and function [13]. Therefore, it is suggesting that variations in the components and structure of the PSD underpin abnormal synaptic plasticity and ultimately lead to neurological disorders after DEHP exposure.

A great deal of attention has been paid to microRNAs (miRNAs) in regulation of synaptic development and plasticity [14-16]. Notably, individual miRNAs are particularly abundant in presynaptic and postsynaptic compartments, regulating synaptic plasticity and activity by modulating the translation of local proteins, including NMDAR, AMPAR, GTPases, cytoskeleton and PSD scaffold proteins [17, 18]. It has been found that miRNAs expression in central nervous system could potentially be utilized as key biomarkers for neurotoxicity induced by environmental neurotoxicants including DEHP [19]. Focusing on miRNA and its targets in PSD may provide insight into new preventive or therapeutic opportunities.

Diet is one of the main environmental factors affecting brain plasticity [20]. Omega-3 fatty acid (Omega-3FA) is classified as essential nutrient since its level depend on dietary intake. Omega-3FA mainly found in foods such as deep-sea fish, marine plants, nuts, and vegetable oils [21]. As a major component in neuronal membrane, Omega-3FA exhibits a wide range of regulatory functions [22], and play critical roles in maintaining brain structure. Omega-3FA deficiency has been related to hippocampal plasticity reduction and memory deficits in rodents, while dietary Omega-3FA supplementation may promote neuroplasticity and improve learning and memory abilities [23, 24]. The increasing evidences support the beneficial effects of an increasing Omega-3FA intake on various neurodegenerative and neurological conditions [25, 26]. In addition, dietary factors have been shown to modify expression of miRNA and their mRNA targets in various neurotoxic processes, including cell cycle regulation, apoptosis, inflammation and neurotransmitter homeostasis as well as pathways in synaptic plasticity [27, 28].

Since Omega-3FA supplementation has been explored against different environmental toxicant induced toxicity and has no known adverse effect [29, 30], our study aimed to explore the protective effect of Omega-3FA against DEHP-induced neurotoxicity by modulating synaptic plasticity via PSD associated miRNAs and their target proteins.

## Materials and methods

Animal handling and procedures used in this study were approved by the Institutional Animal Care and Use Committee of Dalian Medical University. All procedures were carried out under strict accordance with National Institute of Health Guide for Care and Use of Laboratory Animals, and all efforts were made to minimize suffering.

### Animal treatment

Sixty KM mice (Male, 4-week-old) were obtained from the Experimental Animal Center of Dalian Medical University, and randomly divided into three groups, that is, control, DEHP (5mg/kg) (Sigma-Aldrich Corp, USA), and Omega-3 FA (150 mg/kg) (Aladdin, China) + DEHP (5mg/kg) groups (n = 20 in each group). Five mice were housed in each cage with a 12 h dark-light cycle, at 18°C-22°C and 50% humidity and were maintained on a standard diet with water available ad libitum. All mice in each group received treatment simultaneously and continuously for 8 weeks. DEHP was prepared by dissolving in a low volume of corn oil, and the mice were administrated by DEHP or Omega-3 FA via gavage. The body weight of each mouse was weighed every week to adjust the dose of DEHP and Omega-3 FA.

### MWM test

Mice were tested for their learning and memory ability using the Morris water maze (MWM) test. Briefly, the test was composed of the spatial acquisition phase and probe trial [31, 32]. In spatial acquisition phase, a circular transparent escape platform located in the center of the N quadrant was submerged 1 cm below the water surface. Animals were trained four sessions by starting at N, S, E, and W quadrants of the pool. The time spent to reach the platform (escape latency) within 60 s was recorded as acquisition latency.

On the fifth day, a spatial probe test was performed. The platform was taken away, each mouse was started at one point in the S quadrant and allowed to navigate freely in the pool for 60 s. The crossings in the target quadrant (which had a hidden platform previously) were recorded by a smart video tracing system (NoldusEtho Vision system, version 5, Everett, WA, USA).

### Ultrastructure investigation

After 8 weeks treatment, there is not a single animal has died. The mice were anesthetized with sodium pentobarbital via i.p. injection. The brain tissue was removed, and hippocampus samples from each group were collected (−2 mm at the anterior/posterior axis, ±1.8 mm at the lateral/medial axis and −1.5 mm at the dorsal/ventral axis), fixed with 4% glutaraldehyde, cooled and separated into 1 mm^3^ pieces; which were later fixed, dehydrated, soaked, embedded and dual stained as previously described [33]. Finally, ultrathin hippocampus slices (500-700 nm) were prepared and the ultrastructure of the synapses was observed via TEM (JEM-2000EX, Olympus, Tokyo, Japan), and the thickness and width of the PSD was recorded [34].

### miRNA sequencing and bioinformatic evaluation

Total RNA extracted from hippocampus tissue samples were used for library preparations and sequenced on an Illumina Hiseq 2500 platform (Illumina, San Diego, CA, USA). The biological significance of altered miRNA expression is closely related to their gene targets. Potential target genes of the differentially expressed miRNAs were predicted from data in the databases: TargetScan, miRWalk, miRanda, miRNA.org and miRDB, and the final targets were integrated from at least two different programs. The GO functional annotation and KEGG pathway analysis were carried out via DIANA-miRPath v3.0 [35-37].

### Western blot analysis

Western blot analyses were performed to detect the protein expression of mGluR5, Homer1, and NMDAR2B (based on results from miRNA screening). GADPH (housekeeping protein) was used as a control. The total protein was prepared as described previously [38]. Equal amounts of protein were separated via 10% SDS-PAGE gel and electrotransferred to Hybond-P polyvinylidene fluoride membranes (Millipore, France). Then blocked for 1h and incubated overnight at 4°C with primary antibodies against mGluR5, Homer1, and NMDAR2B (1:1000, Abcam, USA). Then membrane was washed and incubated with horseradish peroxidase-conjugated goat anti-mouse IgG (1:7,000) (Sigma) at room temperature for 1 h. Enhanced ECL chemiluminescence was quantified densitometrically by UVP BioSpectrum Multispectral Imaging System (Ultra-Violet Products Ltd, Upland, CA).

### Statistical analysis

Data are presented as the mean ±standard error of mean (SEM), and analyzed using SPSS 23.0 for Windows. Student’s t test or one way ANOVA followed by post hoc tests, were performed where appropriate, and in all instances a *p* value of less than 0.05 was considered to be statistically significant.

## Results

### Omega-3FA improved DEHP-induced impairment of learning and memory

To determine whether Omega-3FA could protect against DEHP-induced impairment of learning and memory, Omega-3FA was administrated in combined DEHP to mice for consecutive 8 weeks. As shown in Fig.1A, MWM test showed that DEHP-induced learning and memory deficits were significantly improved by Omega-3FA. In the spatial acquisition phase, Omega-3FA-treated reduced escape latency compared with DEHP alone group (Fig.1 B). In the probe test, increased crossings over the platform in the target quadrant were showed in Omega-3FA-treated mice compared with DEHP alone group (Fig.1 C).

**Fig 1.**
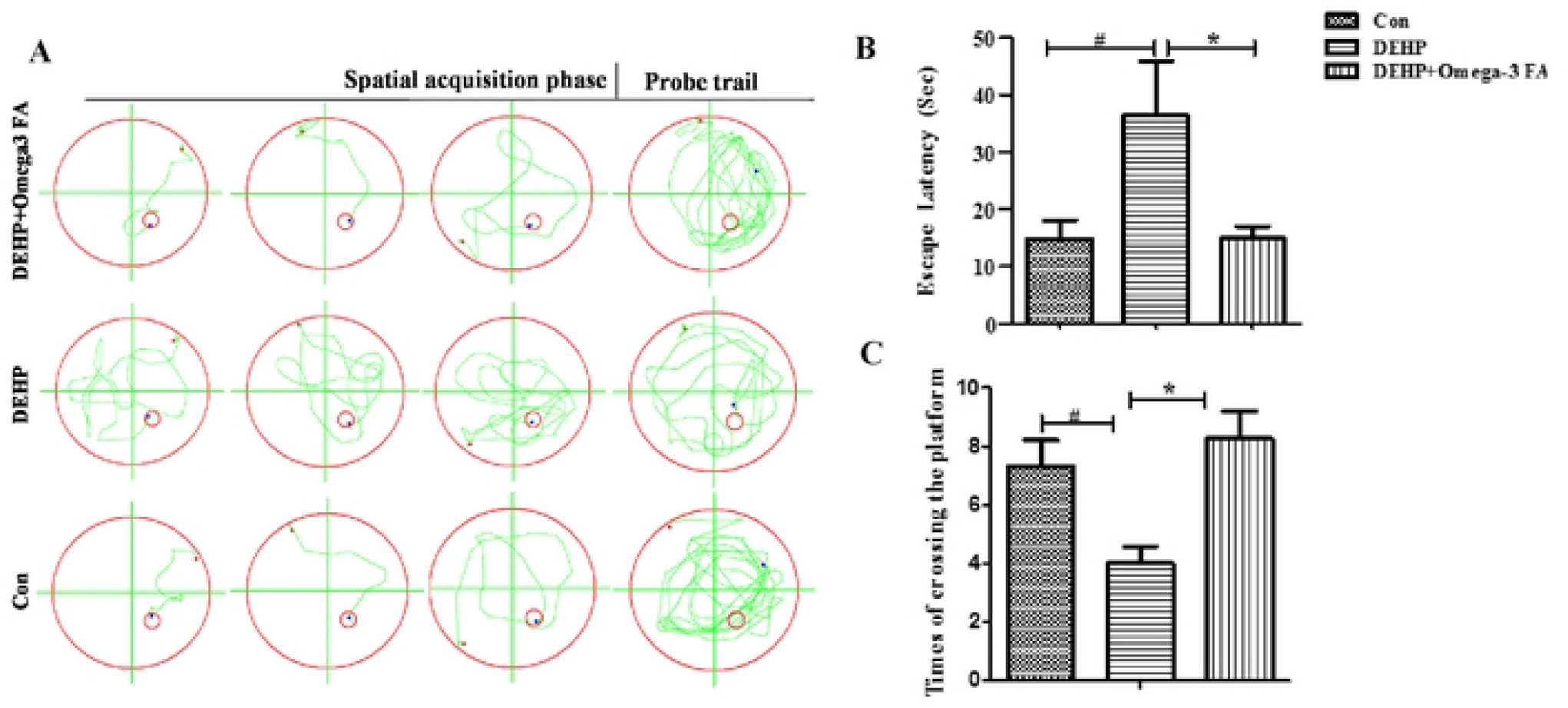
Omega-3FA improved DEHP-induced impairment of learning and memory capacity in mice (n=10-15). Representative swimming routes of each group during the spatial acquisition phase (the 5^th^ day) and probe trial in the Morris water maze (MWM) (A). In spatial acquisition phase, the time spent to find the hidden platform of mice is shown (B). In the probe trial, the total number of crossings the platform in the target quadrant of mice were shown (C). ^#^*p* < 0.05 compared to the control group, \**p* < 0.05 compared to the DEHP group.

### Omega-3FA recovered the synaptic structure in hippocampus of DEHP-treated mice

To investigate whether Omega-3FA could protect against DEHP induced synaptic structural changes, we further examined the structural parameters of the hippocampus synaptic interface.

As shown in Fig.2A, the ultra-structure of hippocampus synapse was obviously recovered in DEHP+Omega-3FA group compared with DEHP alone group. Quantitatively, compared with the control group, decreased PSD thickness and the increased synaptic cleft were detected in the hippocampus synapse of the DEHP group. Treatment with Omega-3FA increased the PSD thickness and decreased the synaptic cleft width significantly compared with the DEHP group (Fig.2B).

**Fig 2.**
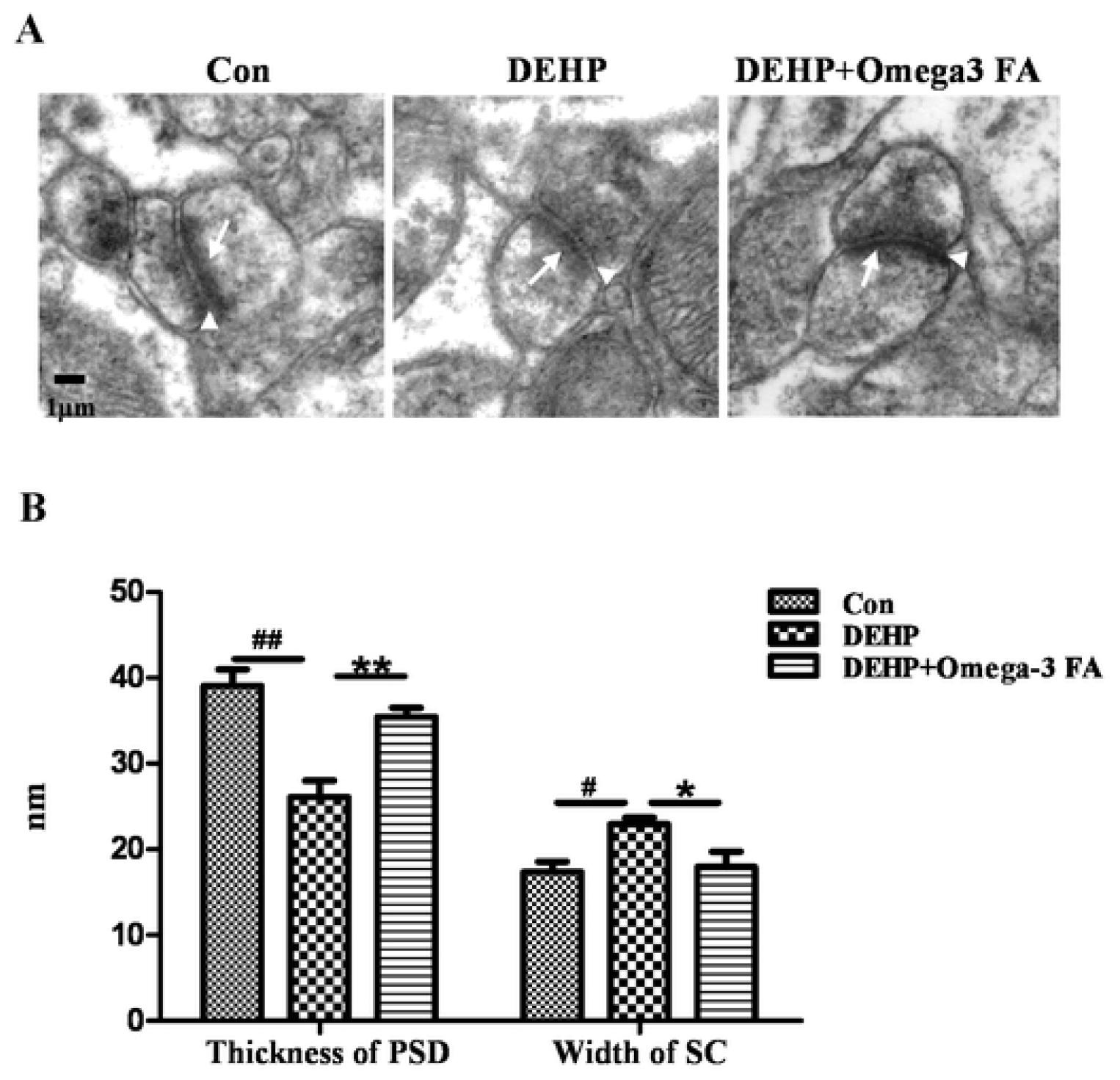
Effects of Omega-3FA on synaptic ultrastructure in hippocampus of mice exposed to DEHP (n=5). Comparison of synaptic structure in the mice hippocampus among groups (A). PSD, postsynaptic density (arrows); SC, synaptic cleft (arrowheads). Quantitatively changes in the PSD thickness and SC width of synaptic in the mice hippocampus among groups (B). ^#^*p* < 0.05, ^##^*p* < 0.01 compared to the control group; \**p* < 0.05, \*\**p* < 0.01 compared to the DEHP group.

### Omega-3FA affects postsynaptic density related miRNAs in alleviating DEHP-induced impairment of learning and memory

Differentially expressed miRNAs were shown in Fig.3A, there were 62 differently expressed miRNAs identified between the DEHP group and control group, as well as there were 89 differently expressed miRNAs between the DEHP+Omega-3FA group and the DEHP group. We found 25 common miRNAs on both lists of differentially expressed miRNAs in DEHP vs CON and DEHP+Omega-3FA vs DEHP groups (Fig.3B). Excluding those with changes in the same direction, we identified twenty miRNAs, upregulated miR-3963, miR-344-3p, miR-335-3p, let-7a-5p, let-7g-5p, miR-145a-5p, miR-122-3p, miR-652-3p, miR-381-3p and miR-26b-5p in DEHP exposure which were downregulated by Omega-3 FA treatment, and downregulated miR-129b-5p, miR-378b, miR-411-3p, miR-338-5p, miR-301a-3p, miR-1195, miR-6238, miR-1970, miR-1969, and miR-690 which were upregulated by Omega-3 FA treatment (Fig.3C).

**Fig 3.**
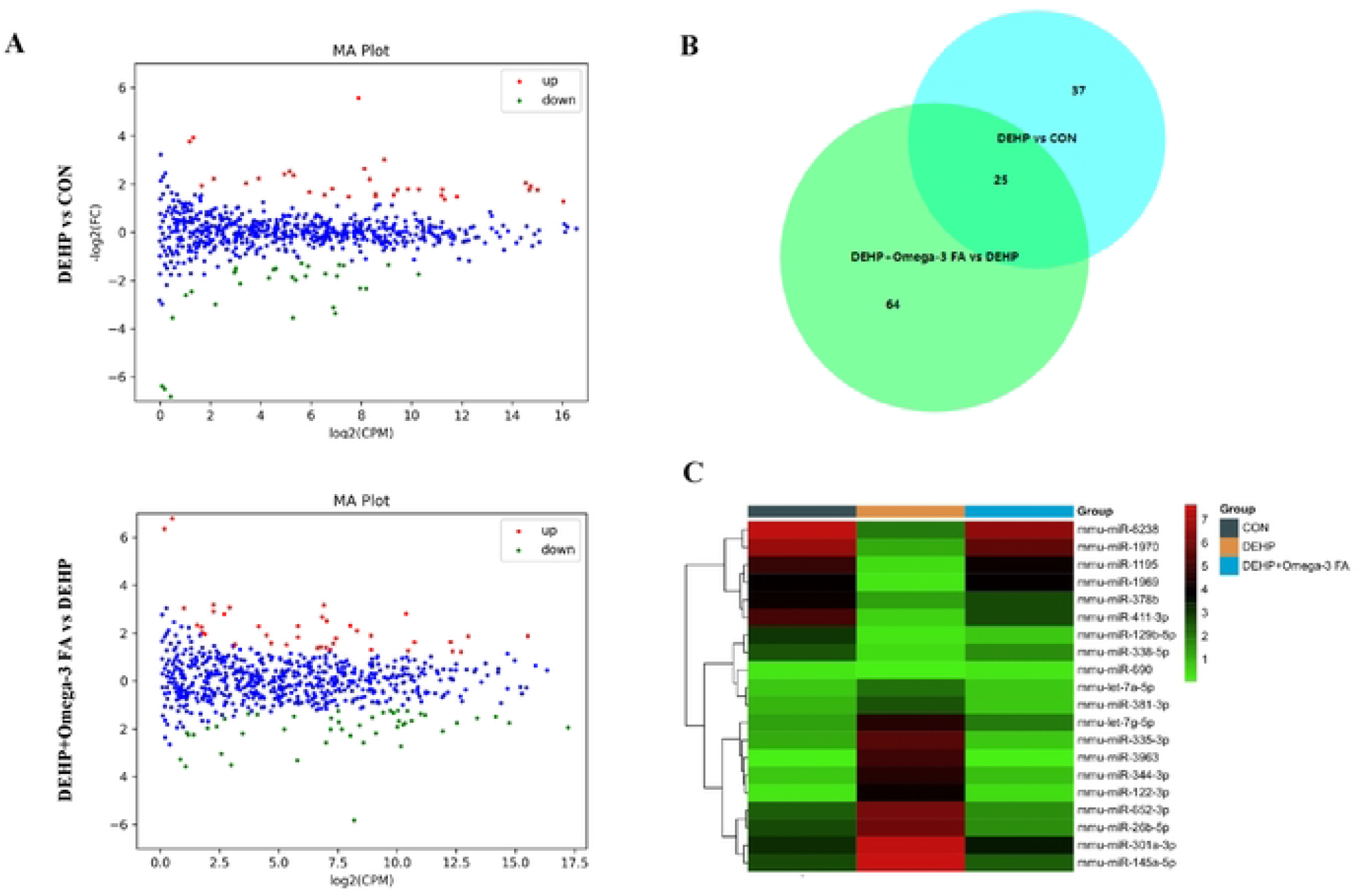
Detail information of differentially expressed miRNAs (n=3). A: Scatter plot of differentially expressed miRNAs in hippocampus. Red and green dots indicate up and downregulation of miRNA, respectively, relative to the control (*P* < 0.05). B:Venn diagram showing the number of significantly modulated miRNAs in DEHP vs control and DEHP+Omega-3 FA vs DEHP. C: Heat map representation of significantly differentially expressed miRNAs (group averages) in control, DEHP and DEHP+Omega-3 FA-treated mice.

After bioinformation analysis of these twenty miRNAs, we focused on the changed miRNAs associated with GO term (GOTERM CC FAT) of “Postsynaptic density” (Table 1). In Table 2, Fourteen differentially expressed miRNAs linked “Postsynaptic density” in DEHP vs CON and DEHP+Omega-3FA vs DEHP groups were listed out, including miR-344-3p, miR-335-3p, let-7a-5p, let-7g-5p, miR-145a-5p, miR-381-3p, miR-26b-5p, miR-129b-5p, miR-378b, miR-338-5p, miR-6238, miR-1970, miR-1969 and miR-690.

**Table 1.** The GO terms significantly changed in cellular component (GOTERM CC FAT) of differentially expressed miRNAs in DEHP vs CON and DEHP+Omega3FA vs DEHP groups.

| **Term** | ***p*-Value** | **Ontology** |
| --- | --- | --- |
| **Synapse** | **5.0E-9** | **Cellular component** |
| **Dendrite** | **1.3E-6** | **Cellular component** |
| **Dendritic spine** | **2.2E-5** | **Cellular component** |
| **Postsynaptic density** | **1.2E-4** | **Cellular component** |
| **Neuronal cell body** | **1.8E-4** | **Cellular component** |
| **Presynaptic membrane** | **3.0E-3** | **Cellular component** |
| **Cytoskeleton** | **9.4E-3** | **Cellular component** |
| **Postsynaptic membrane** | **1.5E-2** | **Cellular component** |

**Table 2.**
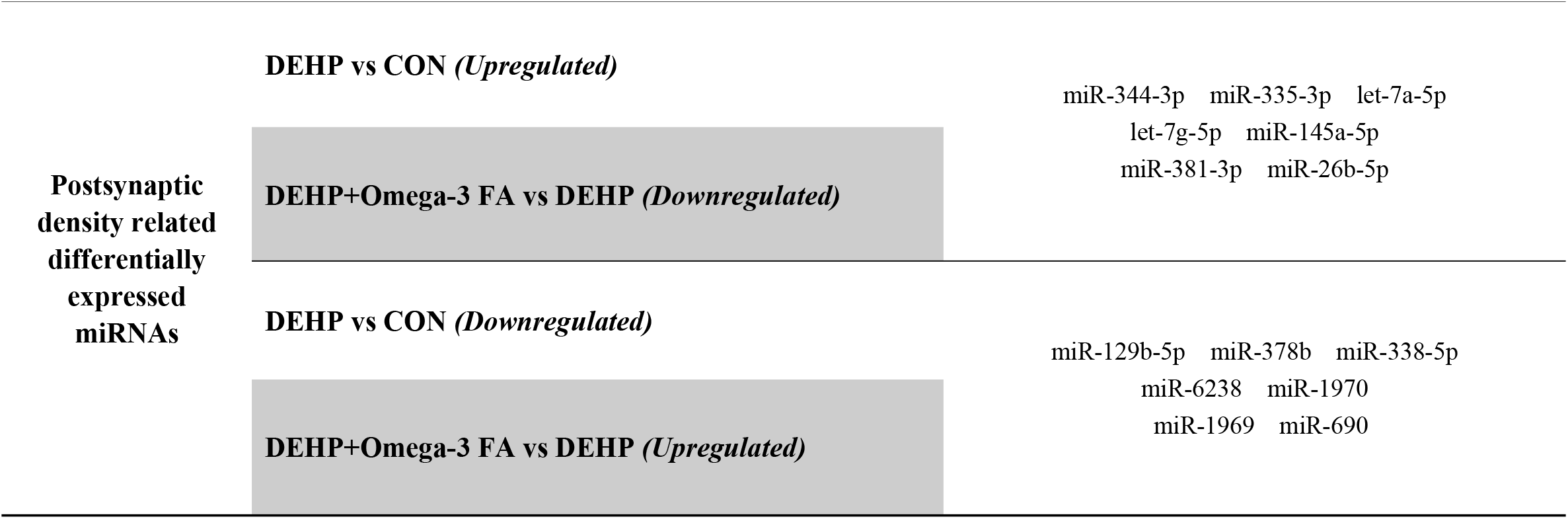
Postsynaptic density related differentially expressed miRNAs in DEHP vs CON and DEHP+Omega-3 FA vs DEHP groups.

### Post synaptic density related miRNAs’ target genes analysis and conformation

The potential mRNA targets of these differentially expressed miRNAs associated with “Postsynaptic density” were further collected using the comparative platforms. Targets calculation was performed by at least 2 different algorithms in case of minimizing the number of putative and maybe false positive targets. Target mRNAs of the PSD related miRNAs were predicted and shown in Fig.4.

**Fig 4.**
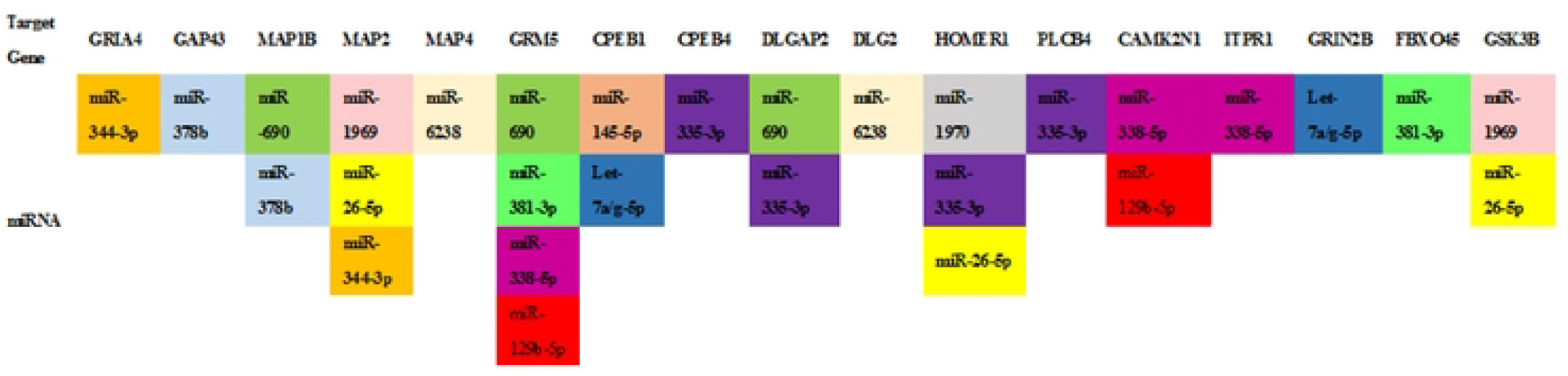
Detail information of postsynaptic density (PSD) regulatory miRNAs’ target genes in PSD (The same miRNAs were filled with one color).

The protein expression levels of the target gene GRM5, HOMER1 and GRIN2B were further investigated, due to the targeted regulation of more than 2 differentially expressed miRNAs. The expression levels of the mGluR5 (GRM5), Homer1 (HOMER1), and NMDAR2B (GRIN2B) proteins in hippocampus are shown in Fig.5. DEHP exposure reduced the expression levels of mGluR5, Homer1, and NMDAR2B protein in hippocampus of mice, which was significantly upregulated by Omega-3FA.

**Fig 5.**
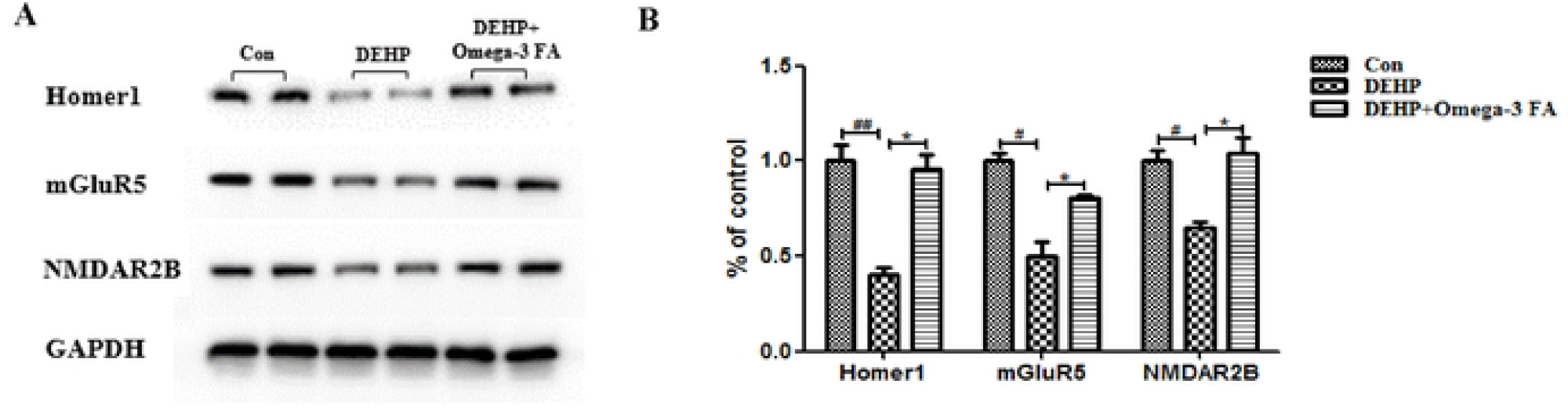
Western blot analysis of the protein levels of Homer1, mGluR5 and NMDAR2B in hippocampus (n=5). ^#^*p* < 0.05, ^##^*p* < 0.01 vs the control group; \**p* < 0.05 vs the DEHP group.

## Discussion

In the present study, we show for the first time that Omega-3FA provides protect effects on DEHP-induced neurotoxicity. Our research has revealed that Omega-3FA supplement could modulate PSD related miRNAs’ expression and improved synaptic structure as well as learning and memory ability, which were weakened by DEHP.

DEHP is a lipophilic compound and well absorbed after oral exposure. In human body, the absorption of DEHP could be 25% [39]. A recent study investigated that DEHP affected the integrity and function of the blood-brain barrier (BBB), which could lead to neurotoxic effects [40]. Omega-3FA is associated with BBB integrity in human [41]. In animal models, Omega-3FA mitigate BBB disruption after brain injury [42]. It is suggesting that the BBB integrity might be protected by Omega-3FA, which may contribute to alleviate partially the impairment of learning and memory induced by DEHP.

An increasing number of studies have suggested that structure synaptic plasticity may present a novel target of Phthalates induced neurotoxicity [43, 44]. Dong et al. recently reported that maternal DEHP exposure affected the synaptic ultrastructure of hippocampus in male offspring [8]. It has been suggested the thickness of the PSD is a crucial indicator of synapses morphology, as the efficiency of synaptic transmission decreases when the PSD becomes thinner [45]. In this study, we found that Omega-3FA administration increased thickness of PSD, which was associated with alleviated cognitive impairment in mice of DEHP exposure.

miRNAs have been identified as potential diagnostic and therapeutic targets in many diseases [46], and recently have emerged as key regulators of neuronal development and synaptic plasticity [47]. One previous study has verified that the levels of various miRNAs are significantly changed in brain tissues after DEHP exposure [19]. It was reported that Omega-3FA affected miRNA expression in hippocampus of animal models [48]. Therefore, we analyzed the effect of Omega-3FA on miRNAs expression after DEHP exposure. Our bioinformatics analysis on differential expression miRNAs showed that “Post synaptic density” was an important cellular component of Omega-3FA protecting against DEHP neurotoxicity. There were fourteen differential expression miRNAs presenting in the “Post synaptic density” term, including: miR-344-3p, miR-335-3p, let-7a-5p, let-7g-5p, miR-145a-5p, miR-381-3p, miR-26b-5p, miR-129b-5p, miR-378b, miR-338-5p, miR-6238, miR-1970, miR-1969 and miR-690. Variation in PSD associated miRNAs expression affect level of synaptic proteins, and consequently modulating synaptic plasticity through the stability and translation of dendritically localized transcripts [49]. As a multimodal hub, PSD is located on the postsynaptic membranes at the synaptic junction, composed of hundreds of different proteins such as scaffold proteins, glutamate receptors, calmodulin binding protein, ion channels, and signaling molecules [50]. These protein components and their interactions may elucidate the mechanism of long-term changes in synaptic plasticity, which underlie learning and memory [51].

In our study, DEHP exposure significantly reduced protein levels of Homer1, mGluR5, NMDAR2B in the hippocampus of mice, which were increased under administration of Omega-3FA, which may associate with improved synaptic structure and behavior. Homer is one of the most widely studied PSD scaffold proteins that closely involved in mechanisms underlying hippocampus-dependent memory processes [52]. Homer proteins are including the transcripts of three mammalian genes, namely Homer 1, Homer 2 and Homer 3, each with many splice variants including the short isoforms Homer1a and Ania3 and long isoforms Homer1b/c and Homer2 and Homer 3[53]. Homer 1-knockout mice exhibit learning deficits [54]. Homer1a, which is produced by an immediate early gene, modulates ligand-independent type I metabotropic glutamate receptor (mGlu5R) signaling [55]. Homer1b anchors mGlu5R to NMDARs and several signaling molecules, such as transient receptor potential ion channels and inositol-1,4,5-trisphosphate receptors [56, 57]. Spatial memory of normal rats can be augmented when Homer 1c is overexpressed in hippocampus [58]. Therefore, Homer 1 protein acts by linking group I mGluRs to NMDARs, as well as by bridging mGluRs with their intracellular downstream effectors [59]. Recent studies have focused on Homer1 exposure to environmental toxicants in rodents [52]. Our result suggested that GRM5/Homer1/NMDAR2 can be affected by DEHP in neuronal tissues. It was found that NMDAR-dependent LTP and mGlu5R-mediated LTD were lacking in adult Omega-3FA-deficient mice [60]. In perinatal mice, dietary Omega-3FA depletion decreased the Homer1 expression, which could be normalized when the Omega-3FA supplemented in time [61]. These results suggest that Omega-3FA protected synaptic plasticity underlying learning and memory ability, related to normalize the expression of GRM5/Homer1/NMDAR2.

In this study, we noticed that Homer1 gene could be targeted by miR-335-3p, miR-1970 and miR-26-5p; mGluR5 gene could be targeted by miR-690, miR-381-3p, miR-338-5p and miR-129-5p; as well as NMDAR2B gene could be targeted by Let-7a/g-5p. It has been confirmed that miR-335 is involved in synaptic plasticity [62]. Shi et al. found that miR-129-5p directly targeting GRM5 [63]. Our results imply GRM5/Homer1/NMDAR2 related miRNAs which associate with synaptic plasticity maybe an important clue for exploring the protective molecular mechanism of Omega-3FA in DEHP induced neurotoxicity.

In addition to the above, our bioinformatics analysis provides more information on possible research clues. NMDAR2B subunit has PDZ-binding domains on their extreme C-termini that are known to interact with the all four members of PSD-95 family and other PDZ proteins [64]. Previous studies have suggested that DEHP alters expression of PSD-95 [12]. Together with our results, it suggested that DEHP affected PSD-95 might be associated with NMDAR2B. MAPs are a group of proteins with either enzymatic or structural activity, which can interact with tubulin polymers. GSK3β is associated with expression of microtubule-protein such as MAP1B, MAP2, APC, CRMP2, and tau [65]. Sun et al suggested that perinatal DEHP exposure affect the GSK-3β expression and increased level of phospho-Tau in hippocampus [66]. In our present study, GSK3β can be regulated by miR-1969 and miR-26-5p; MAP1B, MAP2 and MAP4 which were targeted by miR-1969, miR-26-5p, miR-344-3p, miR-690, miR-378b and miR-6238. Functional networks of miRNAs and their target genes in synapses is complex, as each miRNA may target many mRNAs and each gene could be regulated by multiple miRNAs [67]. The molecular processes in PSD that underlie synaptic plasticity is believed to control the intracellular cross-linking amongst the transduction pathways. Our work reports for the first that miRNAs expression and their targets in PSD can be affected by DEHP in hippocampus.

Taken together, these observations highlight that DEHP exposure could affect the PSD proteins by disturbing miRNAs expression, leading to changes in the morphology of hippocampus postsynaptic density, which may be involved in the mechanisms of DEHP induced learning and memory defects. The protective efficacy of Omega-3FA on neurotoxicity caused by DEHP via modulating expression of PSD related miRNAs and their targets suggests possible targets of treating the neurotoxicity of DEHP as well as Omega-3FA might be a promising candidate for future human study.

## Conflicts of interest

The authors declare that they have no conflict of interest.

## Funding

This work was supported by grants from Undergraduate Innovation Project of Dalian Medical University (No. S202110161005).

## Author Contributions

Cong Zhang conceived and designed the experiments; Muyao Ding, Hongyu Ma and Hui Du performed the experiments; Yinglong Yang and Min Yu analyzed the data; Muyao Ding and Cong Zhang wrote and revised the manuscript.

## Notes

### Competing Interest Statement

The authors have declared no competing interest.

